# Dynamics of primitive streak regression controls the fate of neuro-mesodermal progenitors in the chicken embryo

**DOI:** 10.1101/2020.05.04.077586

**Authors:** Charlene Guillot, Arthur Michaut, Brian Rabe, Olivier Pourquié

## Abstract

In classical descriptions of vertebrate development, the segregation of the three embryonic germ layers is completed by the end of gastrulation. Body formation then proceeds in a head to tail fashion by progressive deposition of lineage committed progenitors during regression of the Primitive Streak (PS) and tail bud (Pasteels, 1937b; Stern, 2004). Identification of Neuro-Mesodermal Progenitors (NMPs) contributing to both musculo-skeletal precursors (paraxial mesoderm) and spinal cord during axis formation by retrospective clonal analysis challenged these notions (Henrique et al., 2015; Tzouanacou et al., 2009). However, in amniotes such as mouse and chicken, the precise identity and localization of these cells has remained unclear despite a wealth of fate mapping analyses of the PS region. Here, we use lineage tracing in the chicken embryo to show that single cells located in the SOX2/T positive anterior PS region contribute to both neural and mesodermal lineages in the trunk and tail, but only express this bipotential fate with some delay. We demonstrate that posterior to anterior gradients of convergence speed and ingression along the PS gradually lead to exhaustion of all mesodermal precursor territories except for NMPs where limited ingression and increased proliferation maintain and amplify this pool of axial progenitors. As a result, most of the remaining mesodermal precursors from the PS in the tail bud are bipotential NMPs. Together, our results provide a novel understanding of the contribution of the PS and tail bud to the formation of the body of amniote embryos.

---

The amniote Primitive Streak (PS), which is the equivalent of the amphibian blastopore, forms at the midline of the embryo during gastrulation. It marks the location where epithelial cells of the epiblast undergo epithelium to mesenchyme conversion and ingress to form the mesoderm and endoderm. At the end of gastrulation, the PS has reached its maximum length and begins to regress, laying in its wake the tissues that will form the embryonic body. When regression is complete, the remnant of the PS morphs into the tail bud, a poorly organized mass of cells located at the posterior end of the embryo, which generates the posterior-most regions of the body. The function of the regressing PS and tail bud in body formation in amniote embryos has been debated since the end of the 19^th^ century (Romanoff, 1960). Early work suggested that the anterior PS and tail bud act as a blastema able to give rise to the paraxial mesoderm and neural tube (Holmdahl, 1925; Romanoff, 1960; Wetzel, 1929). However, subsequent studies from Pasteels argued that the PS of chicken embryos is merely a transit area for already fated progenitors, like the blastopore of lower vertebrates (Pasteels, 1937a). This meant that the end of PS elongation marks the final segregation of the three germ layers, and regression of the PS merely leads to the deposition of cells derived from lineage-committed progenitors organized along the PS and adjacent epiblast. According to this view, the neural plate forms the entire nervous system, including the spinal cord (Fernandez-Garre et al., 2002; Garcia-Martinez et al., 1993). Fate mapping studies in avian and mouse embryos have largely corroborated Pasteel’s hypothesis, showing that at the beginning of PS regression the mesodermal progenitors of the trunk are found in the epiblast along the PS (Kinder et al., 1999; Psychoyos and Stern, 1996; Schoenwolf et al., 1992; Smith et al., 1994; Tam and Beddington, 1987; Wilson and Beddington, 1996). These studies demonstrated that the antero-posterior distribution of mesodermal progenitors in the PS epiblast reflects their future medio-lateral fate. Cells within the node generate the notochord, and the most anterior PS cells produce the paraxial mesoderm, while the territories of the intermediate, lateral plate and extraembryonic mesoderm lie in progressively more posterior regions of the PS.

Pasteel’s dominant view was recently challenged by the discovery of NMPs in mouse (Tzouanacou et al., 2009). NMPs were identified using a technique where clones are randomly labeled with LacZ and the lineage is inferred a posteriori in fixed embryos. This identified bipotent cells generating large clones containing both neural and mesodermal derivatives along the forming body axis. However, this retrospective strategy did not allow to precisely identify and localize these NMPs in the embryo. Grafts of small territories of the epiblast adjacent to the anterior PS in avian or mouse embryos give rise to both neural and mesodermal derivatives (Garcia-Martinez et al., 1993; Wymeersch et al., 2016). In mouse, cells of this region (caudal lateral epiblast and node streak border) co-express the neural marker Sox2 and the mesodermal marker T/Brachyury, and were proposed to include the NMP population (Wymeersch et al., 2016). However, these grafting experiments do not allow to distinguish between a population of bipotential cells and a mixture of precursors committed to each lineage. Moreover, direct reconstruction of the epiblast cell lineage from high-resolution light-sheet imaging of developing mouse embryos did not identify bipotential NMP cells in the anterior PS epiblast but only cells fated to one or the other lineage (McDole et al., 2018). In the chicken embryo, extensive fate mapping of the anterior PS and Hensen’s Node has been performed but did not identify the NMP population either (Brown and Storey, 2000; Iimura et al., 2007; Psychoyos and Stern, 1996; Schoenwolf et al., 1992; Selleck and Stern, 1991). Thus, so far, there is no direct evidence for the existence of NMPs in amniote embryos.

Here, we performed direct lineage analysis of cells in the anterior PS epiblast using two different approaches: a barcoded retroviral library and a Brainbow-derived strategy (Loulier et al., 2014). We show that in the chicken embryo, single SOX2^+^/T^+^ cells of the anterior PS region are NMPs which can contribute to both neural tube and paraxial mesoderm during formation of more posterior regions of the body. As most published fate mappings have been analyzed only after short term and did not study formation of the posterior regions of the embryo, cells of the anterior epiblast were considered monopotent and their bipotentiality went undetected. Thus, our findings reconcile the existence of bipotential NMPs demonstrated in mouse with the large body of amniote fate mappings described in classical developmental biology literature. Moreover, we show that a posterior-to-anterior gradient of cell convergence speed toward the PS coupled to ingression results in the progressive exhaustion of the PS from its posterior end. This leads to the successive disappearance of the territories of the extra-embryonic, lateral plate, intermediate mesoderm and non-NMP paraxial mesoderm precursors. Increased proliferation combined with limited ingression of cells ensure the self-renewal and expansion of the territory of clonally-related bipotential NMPs during body formation which constitutes the last remnant of the PS which contributes to the tail bud. Thus, our work provides a direct demonstration of the existence of NMPs in amniotes and a mechanistic understanding for their sustained contribution to axis elongation.

## Results

### The SOX2/T territory of the anterior PS epiblast appears at the beginning of PS regression

As cells expressing both SOX2 and T represent the most obvious NMP candidates, we first characterized the double positive SOX2/T population in the epiblast during development of the chicken embryo. At stage 4^−^ HH (Hamburger and Hamilton, (Hamburger and Hamilton, 1992)), SOX2 is only expressed in the neural plate in the anterior epiblast and its expression domain progressively expands posteriorly to reach the anterior PS and the adjacent epiblast at stage 4-5HH (Figure 1a, Supplementary Figure 1a-c). SOX2 expression levels show a graded pattern which peaks in the anterior neural plate and decreases in the epiblast lateral to the anterior-most PS (Figure 1a, Supplementary Figure 1d-f). No SOX2 expression is detected in the epiblast adjacent to the posterior PS. At these stages, T expression shows a reverse gradient along the PS, with low levels in the node region and high levels in the posterior end of the PS (Figure 1b, Supplementary Figure 1e-f). At stage 4+HH, these opposing gradients begin to overlap in the epiblast lateral to the anterior PS, defining a SOX2/T territory which forms an inverted V capping the PS (Figure 1c-d, Supplementary Figure 1g-h). At stage 5HH, numerous double-positive SOX2/T cells are found in the epiblast of the anterior PS region, with the underlying mesoderm expressing the paraxial mesoderm-specific marker MSGN1 (Figure 1e-f, level 2-3, Supplementary Figure 1g-h). Posterior to this domain, the epiblast of the PS region expresses T but not SOX2, while the underlying mesoderm maintains a high level of MSGN1 expression (Figure 1e-f, level 4). Posterior to these two domains, in regions of the epiblast corresponding to the prospective intermediate mesoderm/lateral plate and the extraembryonic mesoderm, epiblast cells express T but not SOX2 and no MSGN1 cells are found in the underlying mesoderm (Figure 1e-f, level 5). Therefore, the presumptive paraxial mesoderm territory of the epiblast of the anterior PS can be subdivided into two sub-domains along the AP axis (Figure 1d). An anterior domain where cells co-express SOX2 and T corresponding to the presumptive NMP territory and a posterior Paraxial Mesoderm Precursors (PMP) domain where cells express T but not SOX2. Both domains are characterized by the production of MSGN1-positive paraxial mesoderm precursors.

**Figure 1:**
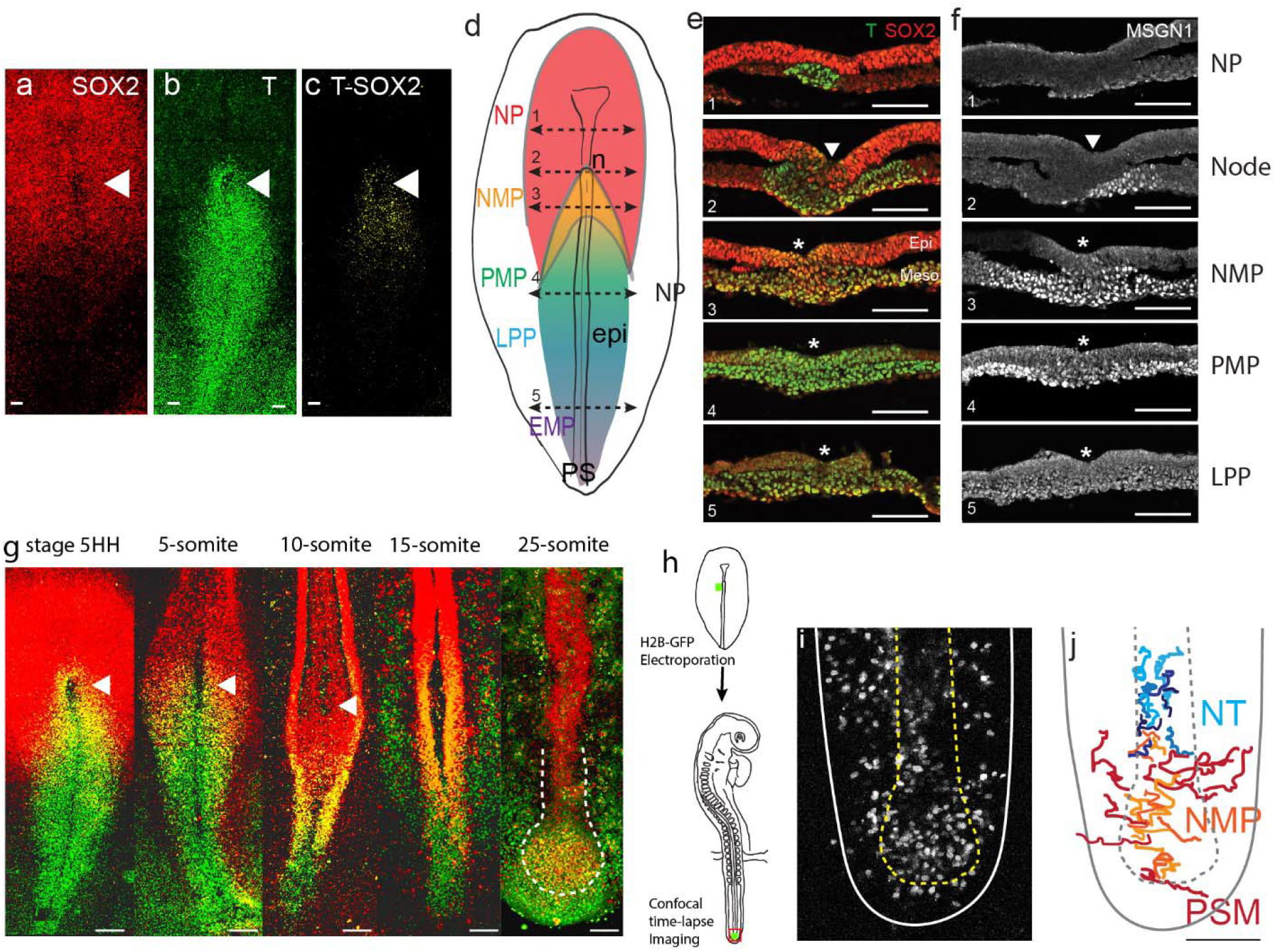
Characterization of the SOX2/T-positive territory of the epiblast of the anterior primitive streak. (a-c) Whole mount embryos and (e-f) transverse cryo-sections showing the immunolocalization of SOX2 (a), T (b), T and SOX2 (c, e), and MSGN1 (f) in chicken embryos at stage 5HH. (d) Schematic representation of the expression of T (blue: high, green: low), SOX2: red and SOX2/T: yellow in a stage 5HH chicken embryo. The level of the tissue sections shown in (e, f) are shown with dashed double arrows labelled from 1 to 5 from anterior to posterior. (n=7 embryos for whole mount; n=3 for cryosections) (g) Maximum intensity projections from confocal images of chicken embryos immuno-stained for T (green) and SOX2 (red) proteins. Double-positive cells are shown in yellow. White hatched line in the 25-somite embryo marks the end of the neural tube (red) and the NMP region (orange) (n=23 embryos analyzed). (h) Diagram summarizing the experimental procedure to label NMP cells using electroporation at stage 5HH of a fluorescent reporter in the epiblast of the anterior PS region (in green) and their analysis after the tailbud stage. (i-j) Fate of descendants of cells of the NMP region electroporated at stage 5 HH with an H2B-RFP plasmid and imaged in time-lapse at the 25-somite stage in the tail bud region. Z-projection from confocal images (i) and tracks (j) of a time-lapse movie showing the movements of the cells in the NMP territory for 10 hours (Supplementary movie 1). Tracks were color-coded a posteriori such as neural, mesodermal and NMP cell trajectories are shown in light blue, red and orange, respectively. NP: Neural Plate, NMP: Neuro-Mesodermal Progenitors, PMP: Presomitic Mesoderm Progenitors, LPP: Lateral Plate Progenitors, EMP: Extraembryonic Mesoderm Progenitors, n: node, epi: epiblast, PS: Primitive Streak. Arrowheads: Hensen’s Node. Asterisk: Primitive streak. (a-d, g-j) Dorsal views. Anterior to the top. Scale bar: 100μm.

### Lineage tracing analysis identifies bipotent cells contributing to the neural tube and paraxial mesoderm in the SOX2/T positive region of the epiblast

In order to explore the fate of the SOX2/T-positive cells of the anterior PS region, we first used a library of barcoded defective retroviruses expressing GFP (Harwell et al., 2015) to infect the epiblast of the anterior PS region at stage 5HH (Figure 2a). Embryos were reincubated for 36 hours and single fluorescent cells were manually harvested from the paraxial mesoderm and neural tube from embryo sections for subsequent barcode analysis. We divided the axis into an anterior territory extending to somite 27 and a posterior territory including the rest of the posterior axis. Somite 27 corresponds to the thoraco-lumbar boundary which marks the transition between primary and secondary neurulation in chicken (Figure 2a) (Le Douarin et al., 1998). We identified 7 clones containing cells expressing the same barcodes (Figure 2b). 4 clones contained cells both in the neural tube and in somites/presomitic mesoderm (PSM) indicating the bipotential nature of the infected cells of the epiblast of the anterior PS region. Descendants of bipotent cells were found in both anterior and posterior regions of the axis (Figure 2b). Three monopotent clones with only neural descendants were also identified (Figure 2b).

**Figure 2:**
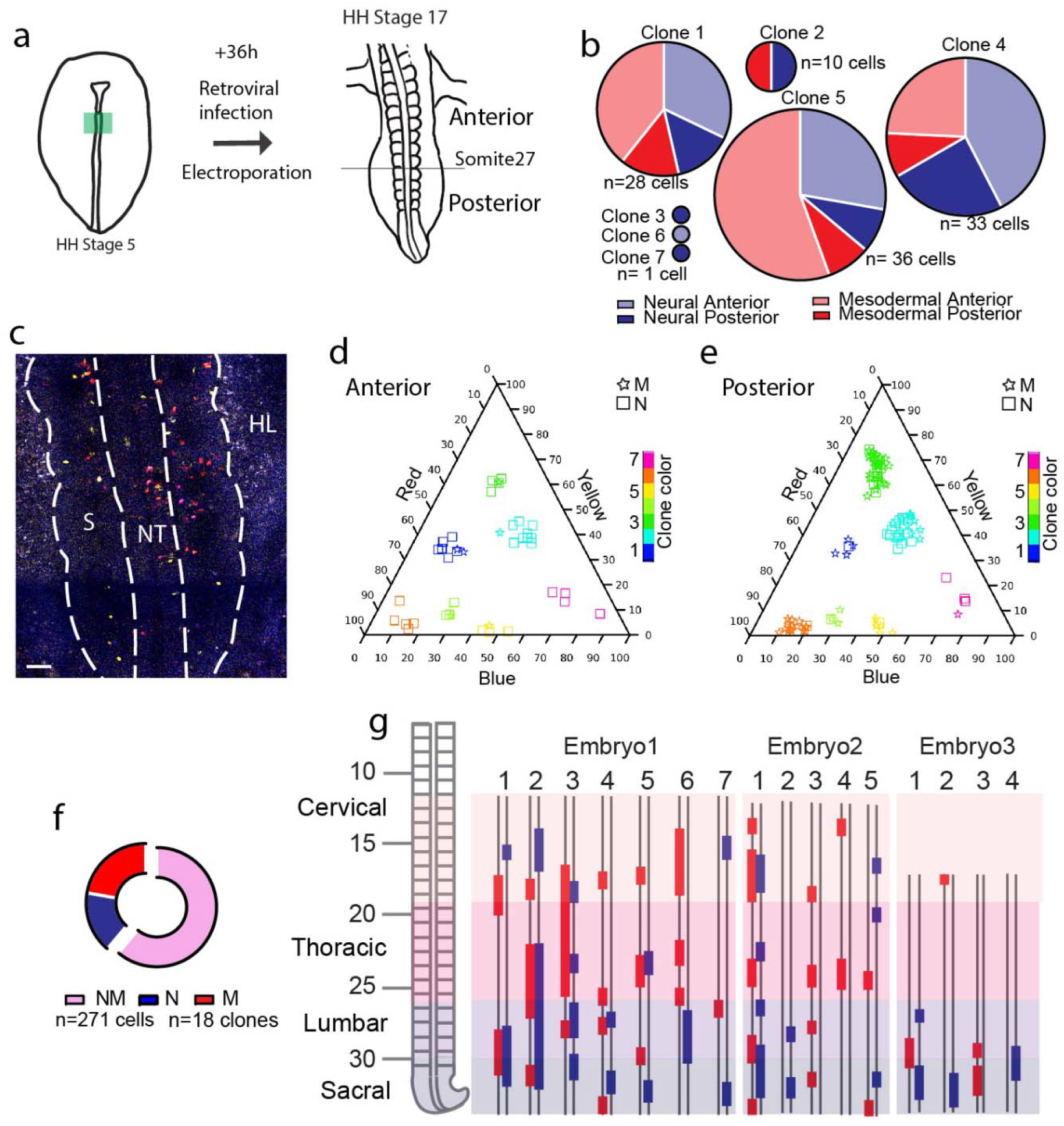
Lineage tracing of SOX2/T double-positive cells shows their contribution to the neural and mesodermal tissues during axis formation. (a) Experimental procedure showing the infected or electroporated region of the epiblast at stage 5 HH (left, green) and the stage at which embryos were harvested for analysis (n=3). (b) Pie graphs showing the distribution of the neural (blue) and mesodermal (red) cells anterior (light) or posterior (dark) to the 27^th^ somite in the seven clones analyzed. Note that some clones exhibit cells only in the anterior (clone 6) or posterior (clones 2,3,7) domains (n=110 cells in 3 embryos). (c) Confocal z-section corresponding to the region of a stage 17 HH embryo shown in (a) and acquired using 3 separated laser paths to retrieve the color codes genetically encoded as described in Loullier et al, 2014. (d-e) Triplot diagrams showing the distribution of descendants of cells labeled with different nucbow combinations in the anterior (d) and posterior (e) region of seven clones in a representative embryo. Each symbol represents a cell identified based on the percentage of red, blue and yellow expressed. The symbols are colored based on their clonal identity. Squares: neural cells, stars: mesodermal cells. (f) Quantification of the different clones: Mesodermal (M, red), Neural (N, blue) and bipotent Neuro-Mesodermal clones (NM, pink). (n= 18 clones, 271 cells in 3 embryos). (g) (Left): Region analyzed showing the different axial levels. (Right): Axial distribution of the clones in 3 embryos. Blue bars: neural cells, red bars: mesodermal cells, double line: AP axis. Dorsal views. Anterior to the top. NT: Neural Tube, S; Somite. HL: Hindlimb. Scale bar: 100μm.

To confirm these observations, we performed lineage tracing of the SOX2/T region of the epiblast using genetic labeling based on the Brainbow-derived MAGIC markers (Loulier et al., 2014). To mark cells, we co-electroporated plasmids expressing a self-excising Cre recombinase and the nucbow transgene together with the Tol II transposase to drive transgene integration. Electroporation of this set of constructs allows to permanently mark cell nuclei with a specific color code generated by the unique combination of different fluorescent proteins triggered by random recombination of the nucbow cassette (Loulier et al., 2014). This color code is then stably transmitted to each daughter cells and can be retrieved by confocal imaging and quantification of the color hues. Using very fine electrodes, we could electroporate as low as 10 epiblast cells of the anterior PS region at stage 5HH (Figure 2a, supplementary Figure 2 a-c). We harvested embryos 36h (stage 17HH) and 56h (stage 20HH) after electroporation (Figure 2a, c, supplementary Figure 3a-b). We identified 58 clones containing a total of 790 cells in the neural tube and paraxial mesoderm in 7 embryos. While we found both monopotent neural and mesodermal clones, the majority of the clones were bi-potent (Figure 2d-g, Supplementary Figure 3d-g). Bipotent clones were found both in the anterior and the posterior region and they often exhibit descendants of only one lineage in some regions and of the other lineage in other regions (figure 2d-e, g, Supplementary Figure 3c, e-g). Thus, our lineage tracing analysis identifies bipotent NMP cells located in the SOX2/T region of the epiblast at stage 5HH. These cells can give rise to neural and paraxial mesoderm progeny in the anterior and posterior regions of the trunk but are not committed to either of these fates.

### Maintenance of the SOX2/T territory during axis formation

We next investigated the fate of the SOX2/T territory during axis formation. During PS regression, i.e. up to the 10-12-somite stage, the SOX2/T cells were maintained in the epiblast lateral to the anterior-most part of the PS, below the Hensen’s node (Arrowhead, Figure 1g). After the 10-somite stage, cells were found in continuity with the posterior-most SOX2 positive/T negative neural tube (Figure 1g). These SOX2/T cells eventually became located in a superficial region of the tail bud at the 25-somite stage where they remain at least until stage 26HH (Olivera-Martinez et al., 2012) (Figure 1g).

In order to analyze the lineage continuity of cells of the anterior PS region, we performed local electroporation of small groups of epiblast cells with a H2B-GFP reporter *in ovo*, targeting the SOX2/T-positive territory of the anterior PS region at stage 5HH (Figure 1h). We next performed confocal live imaging of the electroporated embryos to track the electroporated cells and their progeny at the 25-somite stage (Supplementary movie 1). At this stage, electroporated cells were found in the neural tube, the paraxial mesoderm and the superficial region of the tail bud where the SOX2/T cells were identified (Figure 1g, i, Supplementary Figure 4a-b). Thus, this tail bud territory contains descendants of the SOX2/T cells of the anterior PS epiblast region of stage 5HH embryos. We next performed time-lapse imaging to track fluorescent cells from this superficial SOX2/T positive territory to examine their fate (Figure 1i-j, Supplementary movie 1 and 2). We observed anterior cells undergoing limited movements along the AP axis and moving to join the neural tube (Figure 1i-j, blue tracks, supplementary movie 1 and 2). These cells subsequently acquired the characteristic medio-lateral elongated shape of neural tube cells (Supplementary Figure 4c, Supplementary movie 2). Cells in the middle of the SOX2/T territory moved in the AP direction but undergo very limited medial to lateral movements (Figure 1i-j, orange tracks, Supplementary Figure 4d, Supplementary movie 1 and 2). These cells neither joined the neural tube nor the mesoderm suggesting that they remained in a progenitor state. In contrast, posterior cells undergo dorsal to ventro-lateral cell movements suggesting their ingression in the mesoderm (Figure 1i-j, red tracks, Supplementary Figure 4e, Supplementary movie 1 and 2). Therefore, our data supports lineage continuity and conserved bipotential fate of the SOX2/T territory during axis elongation.

### A posterior to anterior gradient of convergence speed and cell ingression controls the progressive exhaustion of posterior PS precursors and drives the posterior movement of the NMP territory

We next analyzed the cellular dynamics of the SOX2/T territory of the epiblast during PS regression. We performed long-term tracking of epiblast cells from stage 4+HH to 15 somites after nuclear cell labeling using the live marker nuclear red (Supplementary movie 3). We measured the persistence of epiblast cell tracks to localize zones where trajectories end due to cell ingression (Figure 3a; Supplementary Figure 5a, Supplementary movie 4). Posterior cell trajectories are less persistent than anterior ones, indicating that cells in the posterior part of the PS exit the epiblast layer faster than cells of the anterior region. We next measured the angle of epiblast cells trajectories relative to the midline (Figure 3b-c). The tracks of cells in the SOX2/T region show mainly angles between 0° to 45°, indicating limited convergence toward the midline. In contrast, epiblast cells in the posterior half of the PS show angles from 45° to 90° throughout PS regression, suggesting that these cells actively converge toward the midline to join the PS. To measure the speed of cell convergence along the PS, we plotted the instantaneous speed of cells in the Latero-Medial (V_LM_) and Antero-Posterior (V_AP_) directions over time as a function of their position along the PS (Figure 3d). This revealed a low V_LM_ in the anterior PS region compared to the posterior PS. In contrast, the SOX2/T cells show a high V_AP_, similar to that of the node, suggesting that they follow the node posterior movements. V_AP_ decreases progressively in the LP domain to become minimal in the posterior-most region. A similar analysis at a later stage of PS regression (stage 6 HH-5-somites) revealed similar cell dynamics and trajectories for SOX2/T cells maintaining low lateral to medial and high antero-posterior speed (Supplementary Figure 5c-d). Thus, there is a posterior to anterior gradient of convergence speed toward the PS in the epiblast adjacent to the regressing PS (Figure 3e). The decreased track persistence peaking posteriorly suggests the existence of a cell ingression gradient parallel to the convergence speed gradient. To quantify the global dynamics of cell ingression along the PS, we selectively marked the epiblast cells by applying nuclear red dorsally at stage 4+HH. This procedure, when performed *in ovo*, only labels the dorsal epiblast but not the ingressed mesodermal cells or the endoderm. We performed confocal movies from the ventral side of the embryo to measure the increase of mean fluorescence intensity over time along the PS (Figure 3f, Supplementary movie 5). Since only epiblastic cells are marked at the beginning of the movie, the mesodermal layer will progressively acquire new fluorescent cells over time, reflecting the dynamic of cell ingression. We observed a gradual posterior to anterior increase of fluorescence over time in the mesoderm along the PS (Figure 3f). In the anterior-most region, fluorescent cells are observed 120 minutes after posterior cells, indicating a significant delay in the dynamic of ingression of the SOX2/T cells. In addition, we could fit the different curves of intensity measurements along the antero-posterior axis to a linear curve indicating an antero-posterior gradient of cell ingression of epiblastic cells along the PS (Figure 3g). This ingression pattern parallels the distribution of laminin along the PS, which progressively disappears posteriorly (Figure 3h), consistent with a more active ingression behavior. Together, these experiments demonstrate that the epiblast gradient of convergence speed is coupled to graded cell ingression along the PS antero-posterior axis with minimal ingression of the SOX2/T cells.

**Figure 3:**
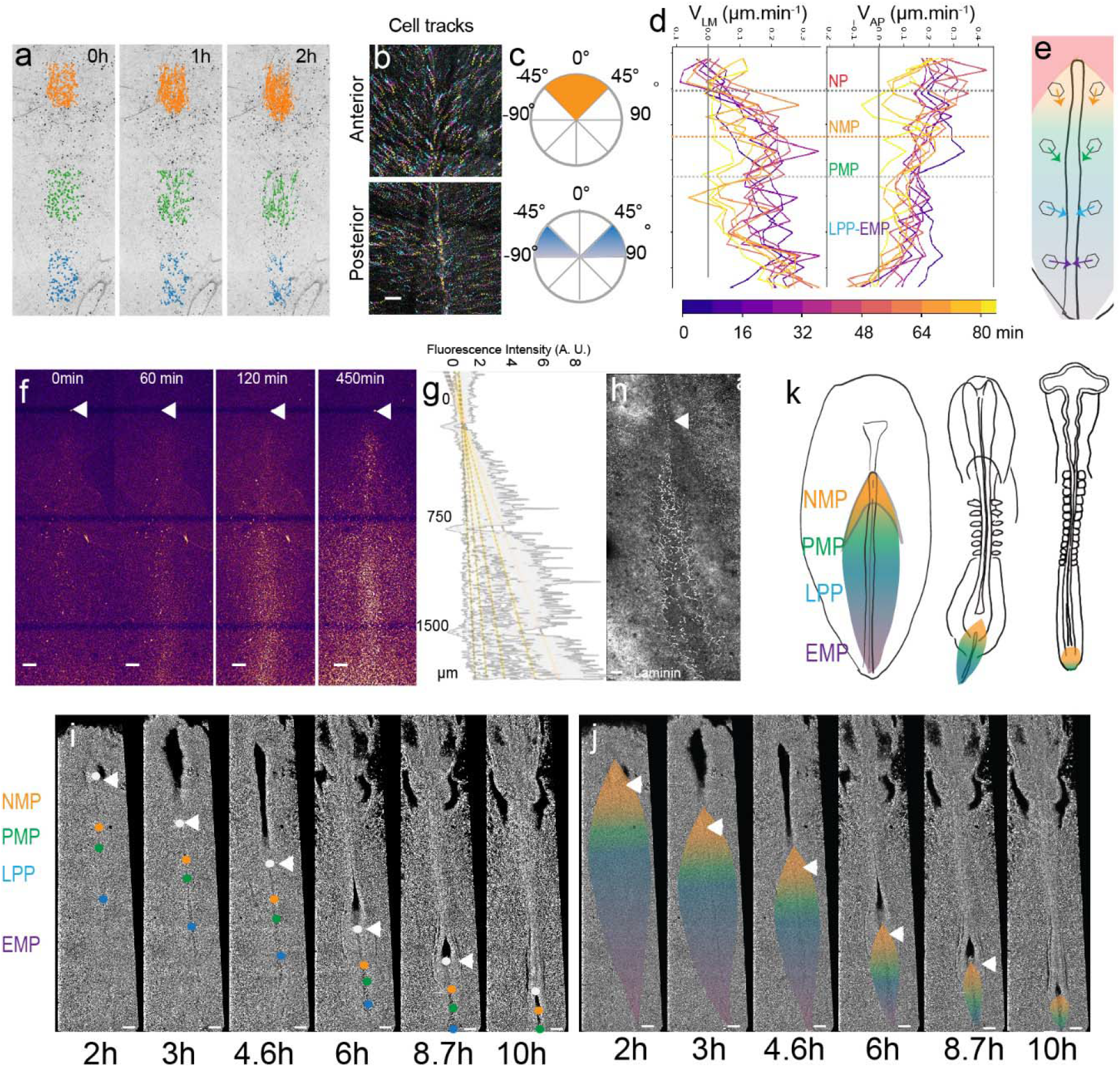
Limited convergence of the NMP territory in the epiblast. (a) Snapshots from a time-lapse movie of a stage 5HH chicken embryo in which the epiblast was labeled with nuclear red (Supplementary movie 4). Tracks of single labeled nuclei are shown at three different PS levels at three different time points (0, 1 and 2h) to illustrate differences in persistence of the cells. (NMP: orange; PMP/LPP: green; EMP: blue). (b) Color-coded time projection showing tracks of epiblast cells at the anterior and posterior PS level after nuclear red labeling in a stage 5HH chicken embryo. The tracks color code represents early timepoints in cyan and later timepoints in yellow. (c) Quantification of the angle with the midline of tracks shown in (b). Top: NMP region, bottom: LPP/EMP region. (d) Mean Lateral to Medial speed (V_LM_) and Anterior to Posterior speed (V_AP_) over time of epiblast cells labeled with nuclear red in stage 5HH chicken embryos. Y axis represents AP position along the embryo. Color code indicates time of measurement since beginning of the movie. (e) Diagram showing the main direction of epiblast cell movements as a function of their AP position in the epiblast. (f) Snapshots from a confocal movie of the PS region of a chicken embryo labeled dorsally at stage 5HH with nuclear red and imaged from the ventral side to show epiblast cells ingression (n=3). (g) Intensity measurement of the nuclear red signal from the ventral side along the PS. Y axis, distance to Hensen’s Node. (h) Whole mount immunohistochemistry with anti-laminin (white) in a stage 5HH chicken embryo. Ventral view. (i-j) Snapshots from a 13h time lapse movie of a chicken embryo starting at stage 5HH. The approximate position of the boundaries is shown by colored dots (i) and the corresponding territories are shown (j) during PS regression. Boundaries between NMP-PMP, PMP-LPP and LPP-EMP are illustrated by orange, green and blue dots (i) and color transition (j) respectively. The initial position of groups of cells marking the boundaries between the different PS territories was identified based on their distance to the Hensen’s node (white arrowhead) as established in our experiments and Psychoyos and Stern (1996). These groups of cells were tracked during PS regression to follow the fate of the different PS territories (n=3 embryos). Dorsal views. (k) Schematics summarizing the dynamics of the NMP territory (orange) during PS regression and PS to tail bud transition. AP: antero-posterior. Neural Plate: NP, red; Neuro-Mesodermal Progenitors: NMP, orange; Presomitic Mesoderm Progenitors: PMP, Green; Lateral Plate Progenitor: LPP, Blue; Extra-embryonic Mesoderm Precursors, EMP, Purple. Arrowhead: Hensen’s Node. Anterior to the top. (n=3 embryos for each experiments). Scale bars: 100μm.

The posterior gradients of convergence speed and ingression are predicted to lead to the progressive disappearance of the precursor territories of the PS in a posterior to anterior order. To test this hypothesis, we generated time-lapse movies of GFP-expressing transgenic chicken embryos from stage 5HH to 10-somite. We tracked specific positions along the PS approximately corresponding to the boundaries between the NMP and paraxial mesoderm progenitors (PMP), the PMP and lateral plate progenitors (LPP) and LPP and extraembryonic mesoderm (EMP). To do so, we manually tracked from stage 5HH the trajectories of cells in small regions located at distances of 500 μm, 700 μm and 1000 μm from the node (Figure 3i-j, Supplementary movie 6). We observed a faster reduction of the posterior PS domains, with the extraembryonic territory disappearing first, followed by the LP territory whose ingression is completed at the 10-somite stage (figure 3j-k) (Moreau et al., 2019; Spratt Jr., 1947). After this stage, most T-positive progenitors of the superficial layer of the tail bud also express SOX2 suggesting that they correspond to the remnant of the epiblast flanking the anterior PS and remain the only axial progenitors left in the tail bud (Figure 3k).

### An anterior to posterior gradient of proliferation counteracts ingression in the SOX2/T territory

We noted that the number of SOX2/T cells gradually increases from stage 4+HH to reach a peak around 30 somites (Figure 4a). As limited convergence and ingression is observed in the NMP territory, this increase is most likely explained by cell proliferation. During PS regression, we observed a higher number of phospho-Histone 3 (pH3)-positive cells relative to the total number of cells (mitotic index) in the anterior region of the PS compared to more posterior ones (Figure 4b-c). We performed confocal live imaging of fluorescent H2B-cherry quail embryos and observed more dividing cells in the anterior PS compared to more posterior regions at the same developmental stage (Figure 4d). This confirmed the existence of a higher mitotic index in the SOX2/T territory compared to more posterior regions of the PS (Stern, 1979) (Figure 4b-d). We next manually tracked individual electroporated cells and measured the time spent in the PS prior to cell ingression (Figure 4e-f). All the cells tracked in the mid PS region spent from 1 to 3 h in the PS before ingressing. In contrast, only 35% of the tracked cells of the SOX2/T region show such fast ingression dynamics (within 1-3 h) whereas 65% remain in the PS for more than 7 hours (Figure 4f).

**Figure 4:**
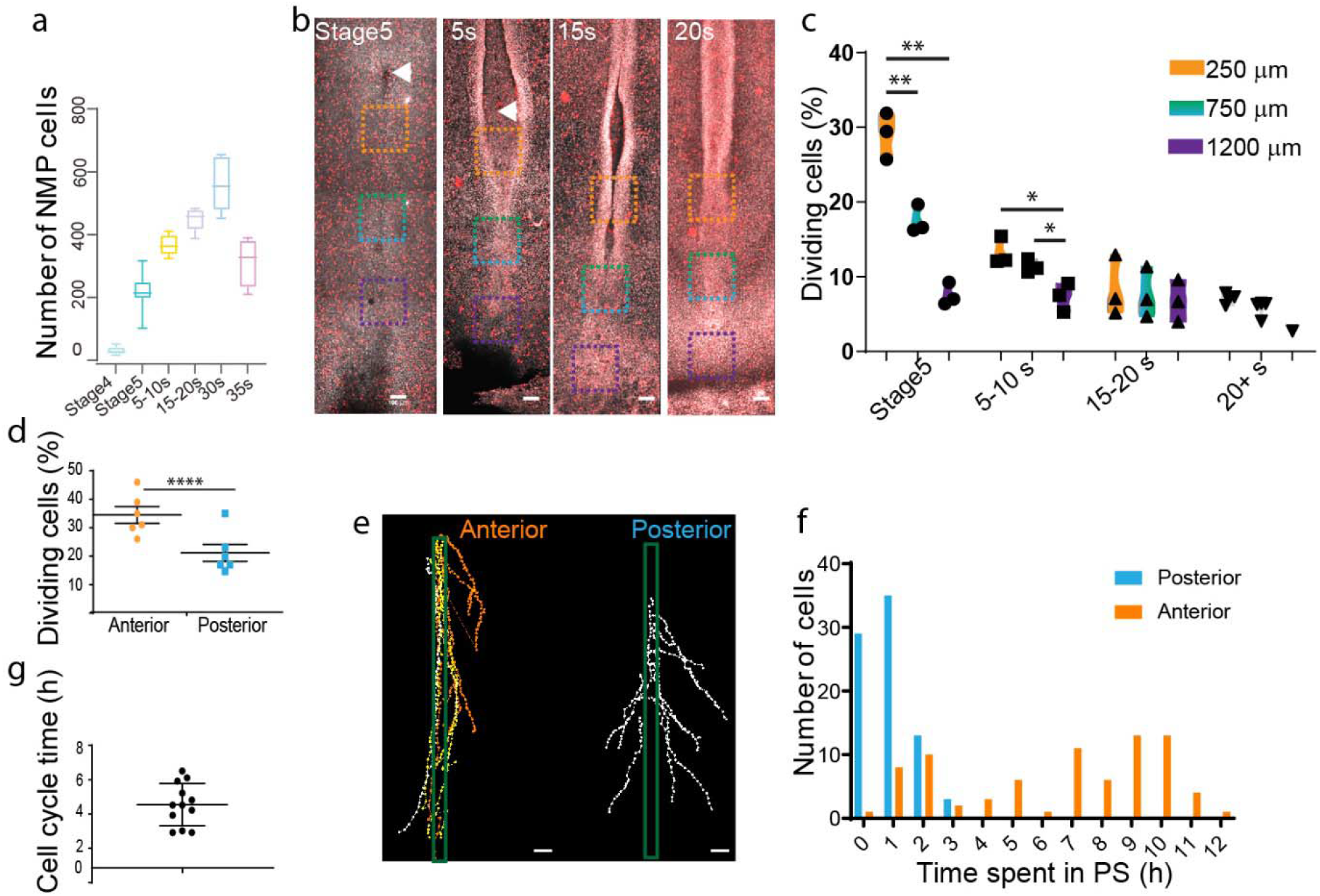
An anterior to posterior gradient of proliferation counteracts ingression in the SOX2/T territory. (a) Quantification of the number of SOX2/T double-positive cells in chicken embryos from stage 4 HH to 35 somites (n=38 embryos). (b) Snapshots of the posterior region of chicken embryos from stage 5HH to 20 somites stained in whole-mount with an anti-phosphorylated Histone H3 (pH3) antibody. (c) quantifications of the mitotic index along the PS in the boxes shown in (b). Orange box: 250 μm from node, Grey box: 750 μm from node, Blue box: 1200 μm from node. (n=13 embryos) Unpaired t-test; **; p=0.0017 and 0.0022; * p=0.0187, p=0.0278. (d) Quantification of the number of dividing cells in H2B-cherry transgenic quails at stage 4+/5 HH in the anterior and posterior PS. (n=6) Paired t test; ***, p= 0.0001. (e) Tracks and (f) quantification of trajectories of the NMP (orange) and LPP (grey) cells during PS regression. (n= 159 cells, 80 posterior, 79 anterior in 7 embryos). (g) Quantification of the time interval between 2 rounds of division in cells of the NMP region measured in time lapse movies. (n=12 inter-division events in 4 embryos). Arrowhead: Hensen’s Node. Dorsal views. Anterior to the top. Scale bar: 100μm.

The number of SOX2/T cells increases from 50 to 550 cells in around 40 hours (Figure 4a). Knowing the percentage of ingressing cells in the anterior PS region (35%), we can predict the evolution of the cell population using a geometric series formula: U_n_=q^n^xU_0_ classically used in analysis of population dynamics. Here n is the number of cell divisions, q is the doubling parameter of the non-ingressed population, (here 2*0.65=1.3) and U_0_ the initial population, i.e. 50 cells. Analysis of the cell cycle length of cells during formation of the posterior tissues in the chicken embryo showed that PSM cells divide every 8 to 10 hours (Benazeraf et al., 2017), thus 4 to 5 times in 40 hours. Using the formula above, the total number of cells after 4 or 5 cell cycles would be U_4_=142 or U_5_=185 cells, i.e. far below the observed number of 550 cells. This suggests that these cells might divide faster than their descendants in the PSM. To obtain a population of 550 cells, the model predicts n=9 or 10 cell cycles (U_9_=530, U_10_=689), suggesting a cell cycle time around 4h. We manually tracked individual dividing cells and their daughter cells in the SOX2/T region over 10 hours to determine the time between two divisions. We identified symmetric cell divisions where the two daughter cells remain in the epiblast of the SOX2/T territory after cell division, consistent with self-renewal of this population. The cell cycle time for such symmetric divisions is around 4.5 hours (Figure 4g, Supplementary Figure 6a, Supplementary movie 7). This cell cycle length is in agreement with the number of divisions predicted by the model above. Other symmetric cell divisions gave rise to two daughter cells entering the mesoderm (Supplementary Figure 6b). We also observed asymmetric cell divisions where one of the daughter cells ingresses after cell division, thus suggesting a specification of one of the daughter cells to a mesodermal fate (Supplementary Figure 6b). Thus, we show that the SOX2/T cells exhibit rapid cell divisions together with limited cell ingression, allowing their self-renewal and amplification during formation of the posterior body.

## Discussion

Our work shows that while most of the PS behaves as a transit zone for committed progenitors as suggested by Pasteels (Pasteels, 1937b), its anterior-most region behaves more like a blastema, containing bipotential NMP cells as predicted by Wetzel (Romanoff, 1960; Wetzel, 1929). In chicken embryos, SOX2/T positive cells are first found in the epiblast adjacent to the anterior PS and Hensen’s Node region at the beginning of PS regression. This domain appears at a similar stage of mouse development and occupies a position similar to the SOX2/T domain of the Node-streak border and the caudal lateral epiblast of mouse embryos (Wymeersch et al., 2016). In both mouse and chicken embryos, this domain contains cells fated to give rise to both neural and mesodermal derivatives, suggesting that they are functionally equivalent (Garcia-Martinez et al., 1993; Wymeersch et al., 2016). We performed lineage tracing using a barcoded retroviral library and brainbow-derived MAGIC markers (Loulier et al., 2014) to show that single cells of the SOX2/T region contribute to the neural tube and paraxial mesoderm along the trunk axis in chicken embryos. We further demonstrate clonal continuity between early NMPs of the PS and late ones in the tail bud. A significant contribution of cells of the SOX2/T territory to both neural and mesodermal lineages has not been documented in most fate mapping studies of this region in chicken embryos (Brown and Storey, 2000; Fernandez-Garre et al., 2002; Henrique et al., 1997; Iimura et al., 2007; Psychoyos and Stern, 1996; Schoenwolf et al., 1992; Selleck and Stern, 1991). Compared to these studies, we analyzed our lineage tracing experiments at significantly later stages (stage 17-20HH instead of stage 10-14HH). Thus, while these fate maps indicate that cells of the anterior PS epiblast initially produce descendants either in the neural tube or in the mesoderm, our results indicate that NMPs give rise to both neural tube and mesoderm only later, in more posterior regions of the body. These observations are consistent with recent grafts of the epiblast territory in 6-somite chicken embryos showing that the territory first produces neural and then both mesodermal and neural derivatives (Kawachi et al., 2020). Bipotential cells with a neural and mesodermal fate have so far only been reported in zebrafish where they segregate during gastrulation and contribute to the most posterior part of the axis (Attardi et al., 2018). In zebrafish however, only monopotent cells were found in the tail bud (Attardi et al., 2018; Kanki and Ho, 1997), while that this is not the case in chicken (this report) and mouse (Tzouanacou et al., 2009).

As reported for mouse embryos (Wymeersch et al., 2016), we observe an increase in SOX2/T cell numbers during axis elongation, suggesting that these cells can self-renew while giving rise to progeny in the paraxial mesoderm and in the neural tube. Labeling experiments in mouse and chicken embryos have identified a population of epiblast cells in the region of the anterior PS and Hensen’s node which behave as stem cells, giving rise to descendants in the paraxial mesoderm while being able to self-renew (Cambray and Wilson, 2002, 2007; Iimura et al., 2007; McGrew et al., 2008; Nicolas et al., 1996; Selleck and Stern, 1991; Wilson et al., 2009). These cells were proposed to contribute mostly to medial somites while lateral somitic cells are derived from more posterior areas of the PS (Iimura et al., 2007; Selleck and Stern, 1991). The anterior SOX2/T territory encompasses the Hensen’s Node and the epiblast adjacent to the anterior PS and approximately corresponds to the territory containing the stem cells fated to give rise to medial somites. The epiblast territory immediately posterior to the SOX2/T territory does not express SOX2 and exhibits many cells positive for the paraxial mesoderm specific marker MSGN1. This territory likely corresponds to the prospective territory of lateral somites which does not show a stem cell behavior, giving rise to descendants spanning only ~5-7 segments (Iimura et al., 2007; Psychoyos and Stern, 1996). Our data suggest that this territory becomes largely exhausted at the end of PS regression resulting in posterior somites to derive mostly from the NMPs. This is consistent with observations in chicken and mouse demonstrating that the selective contribution of different PS territories to medial or lateral somitic territories only applies to anterior somites (Cambray and Wilson, 2007; Psychoyos and Stern, 1996).

We observed a posterior to anterior gradient of convergence in the epiblast associated to a parallel gradient of ingression in the PS. A posterior to anterior gradient of cell motility of ingressed mesodermal cells has been documented along the regressing PS (Zamir et al., 2006). These graded movements in the mesoderm are largely controled by a posterior to anterior gradient of Fgf8 established in the PS and acting as a repellent on newly ingressed mesodermal cells at this stage (Yang et al., 2002). This cellular dynamics suggests that the movement away from the posterior PS could act as a sink for the epiblastic territories generating these progenitors. These cell movements in the epiblast and the ingressing mesoderm could explain the progressive exhaustion of the PS from its posterior end first described for the extraembryonic territory (Spratt Jr., 1947). Importantly, our data show that the precursor territories along the PS do not disappear at the same rate. We observe a sequential posterior to anterior exhaustion of the territories of the extraembryonic mesoderm, the lateral plate and the SOX2 negative paraxial mesoderm progenitors (Figure 3k). Combined to an increased proliferation in the NMP region, this would explain why the SOX2/T territory eventually remains as the major remnant of the PS in the tail bud after PS regression. Thus, the tail bud is composed of a mosaic of monopotent territories such as the precursors of the notochord, or the hindgut, and the multipotent NMPs, which give rise to descendants in the ectoderm and mesoderm. This likely explains why the tail bud appears as a site where gastrulation movements are still ongoing but also shows blastema characteristics (Catala et al., 1995; Davis and Kirschner, 2000; Gont et al., 1993; Holmdahl, 1925).

The difficulty to observe NMPs *in vivo* contrasts with the ease with which NMP-like cells can be obtained *in vitro* from mouse or human pluripotent stem cells. NMPs have been generated in 2D cultures by activation of Wnt signaling at the epiblast-like stage (Diaz-Cuadros et al., 2020; Edri et al., 2019a; Edri et al., 2019b; Gouti et al., 2017; Gouti et al., 2014; Henrique et al., 2015; Turner et al., 2014). Analysis of 3D cultures such as gastruloids induced *in vitro* from mouse ES cells also shows that most of the cells forming these structures belong to the neural tube and paraxial mesoderm lineage, suggesting that they are largely derived from an initial NMP population (Beccari et al., 2018; Faustino Martins et al., 2020; van den Brink et al., 2020). NMP-like cells generated *in vitro* express both SOX2 and T and single cell RNA-sequencing demonstrated that they form a very homogeneous population with characteristics similar to the endogenous SOX2/T cells (Diaz-Cuadros et al., 2020; Gouti et al., 2017). As reported *in vivo* in fish, mouse and chicken embryos, inhibiting Wnt signaling can bias NMP cells toward the neural fate at the expense of the mesodermal fate (Chapman and Papaioannou, 1998; Diaz-Cuadros et al., 2020; Gouti et al., 2017; Gouti et al., 2014; Martin and Kimelman, 2012; Oginuma et al., 2020; Oginuma et al., 2017; Yamaguchi et al., 1999). Here, we show that NMPs can contribute to a single type of derivative (neural or mesodermal) for several segments followed by another type in the next segments, as reported in mouse embryos (Tzouanacou et al., 2009). This argues for a striking plasticity of NMPs and rules out simple models of asymmetric divisions for the generation of neural and mesodermal descendants. Plasticity of the NMPs is supported by heterotopic grafts of the NMP epiblast territory into territories fated to become neural or mesodermal, which resulted in the donor cells to adopt the fate of their new territory in chicken or mouse embryos (Garcia-Martinez et al., 1997; McGrew et al., 2008; Wymeersch et al., 2016). This suggests that the NMP population remains uncommitted toward either lineage, with its descendants acquiring their identity only after entering the territory of the paraxial mesoderm or the neural tube.

## ACKNOWLEDGMENTS

We thank members of the Pourquié lab, Domingos Henrique, Connie Cepko, Denis Duboule and Cliff Tabin for discussions and critical reading of the manuscript. We also thank Jean Livet for the MAGIC markers plasmids. Research in the Pourquié lab was funded by a grant from the National Institute of Health (RO1HD097068-02) and EMBO ALTF 406-2015 to C.G.

## AUTHOR CONTRIBUTIONS

C.G. designed, performed and analyzed biological experiments with O.P. A.M. wrote the code and analyzed the cell trajectories and their speed. B. R. constructed the viral library and prepared the viral solutions. C.G. and O.P. wrote the manuscript; O.P. supervised the project. All authors discussed and agreed on the results and commented on the manuscript.

---

**Supplementary Figure 1 :**
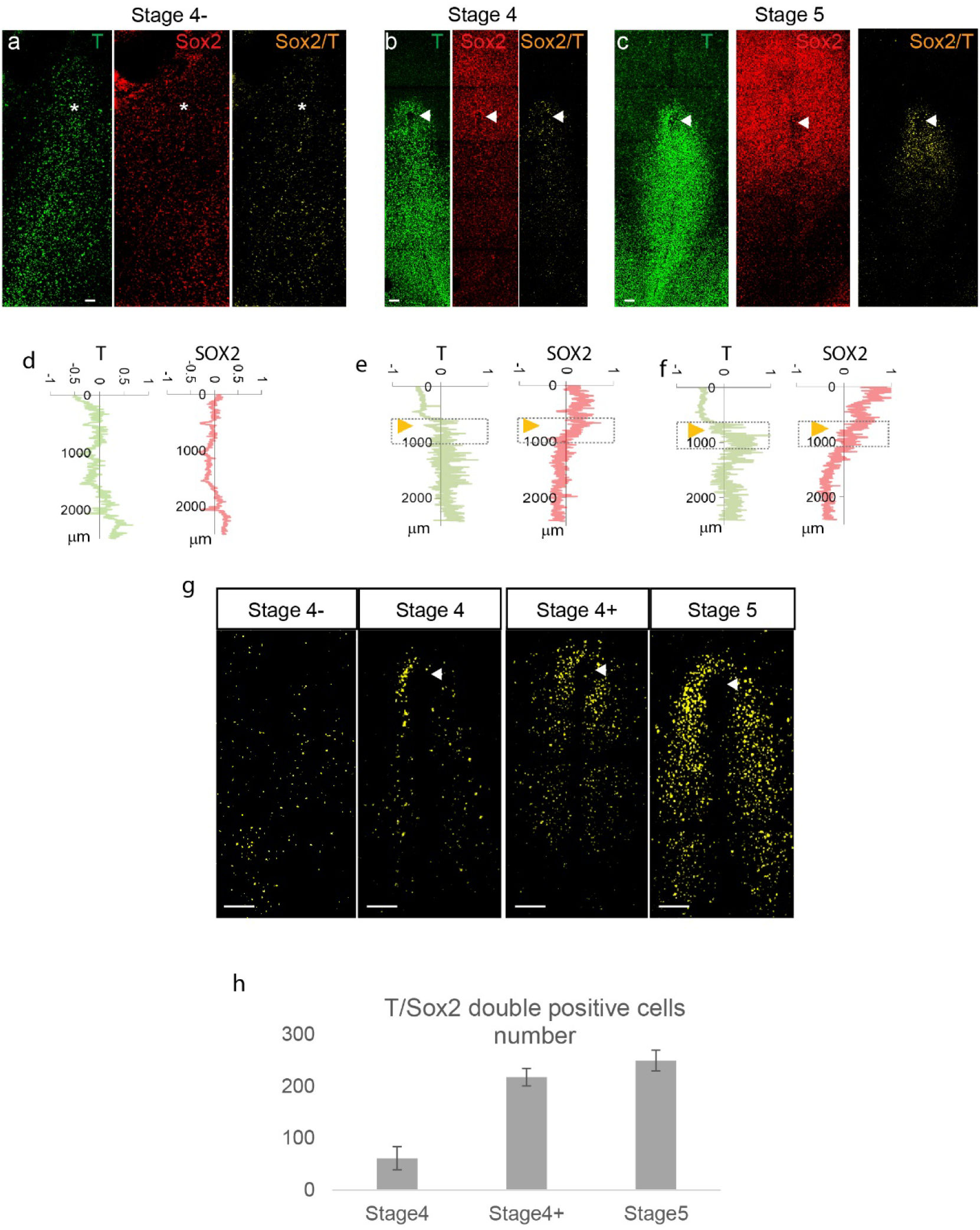
Onset of SOX2/T expression cells in chicken. (a-c) Representative maximum z projections of confocal images of T and SOX2 whole mount immunohistochemistry from stage 4- to stage 5HH. (d-f) Representative line profile intensities showing the quantification of the fluorescence intensity of the SOX2 (red) and T (green) staining along the anteroposterior axis of the embryos (y axis). 0 marks the anterior neural Orange arrowheads: position of Hensen’s Node representative of the anterior border of the PS. (g) Maximum z-projection of embryos stained in whole mount with SOX2/T antibodies showing the localization and the number of the SOX2/T double-positive cells (in yellow) in the anterior PS region. Single positive cells are not shown. (h) Quantification of the number of SOX2/T double positive cells. Dorsal views, anterior to the top. Asterisk marks the tip of PS. White arrowheads: Hensen’s Node. (n= 18; 7 embryos at stage 4HH, 4 at stage 4+HH, 7 at stage 5HH). Scale bar: 100μm.

**Supplementary Figure 2:**
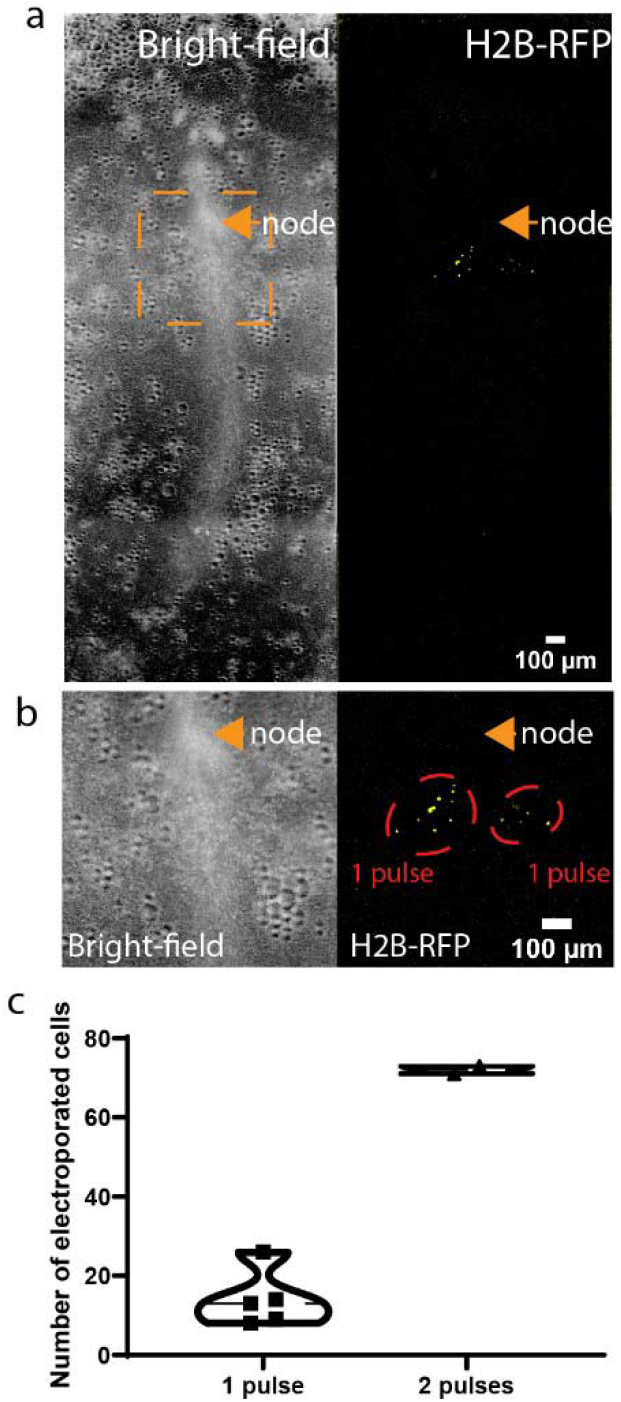
Quantification of the number of epiblast cells electroporated. (a) Brightfield (left) and fluorescence (right) images of a stage 5HH chicken embryo 4h after 1 pulse of electroporation on each side of the anterior PS with an H2B-RFP plasmid. The electroporated areas are within the orange box. (b) Higher magnification of the orange box shown in (a). Fluorescent nuclei in electroporated cells are shown in yellow and the electroporated areas are circled in red. (c) Violin plot showing the quantification of the number of fluorescent nuclei 4h after one pulse or two successive pulses of electroporation at the same location (n= 4 embryos).

**Supplementary Figure 3 :**
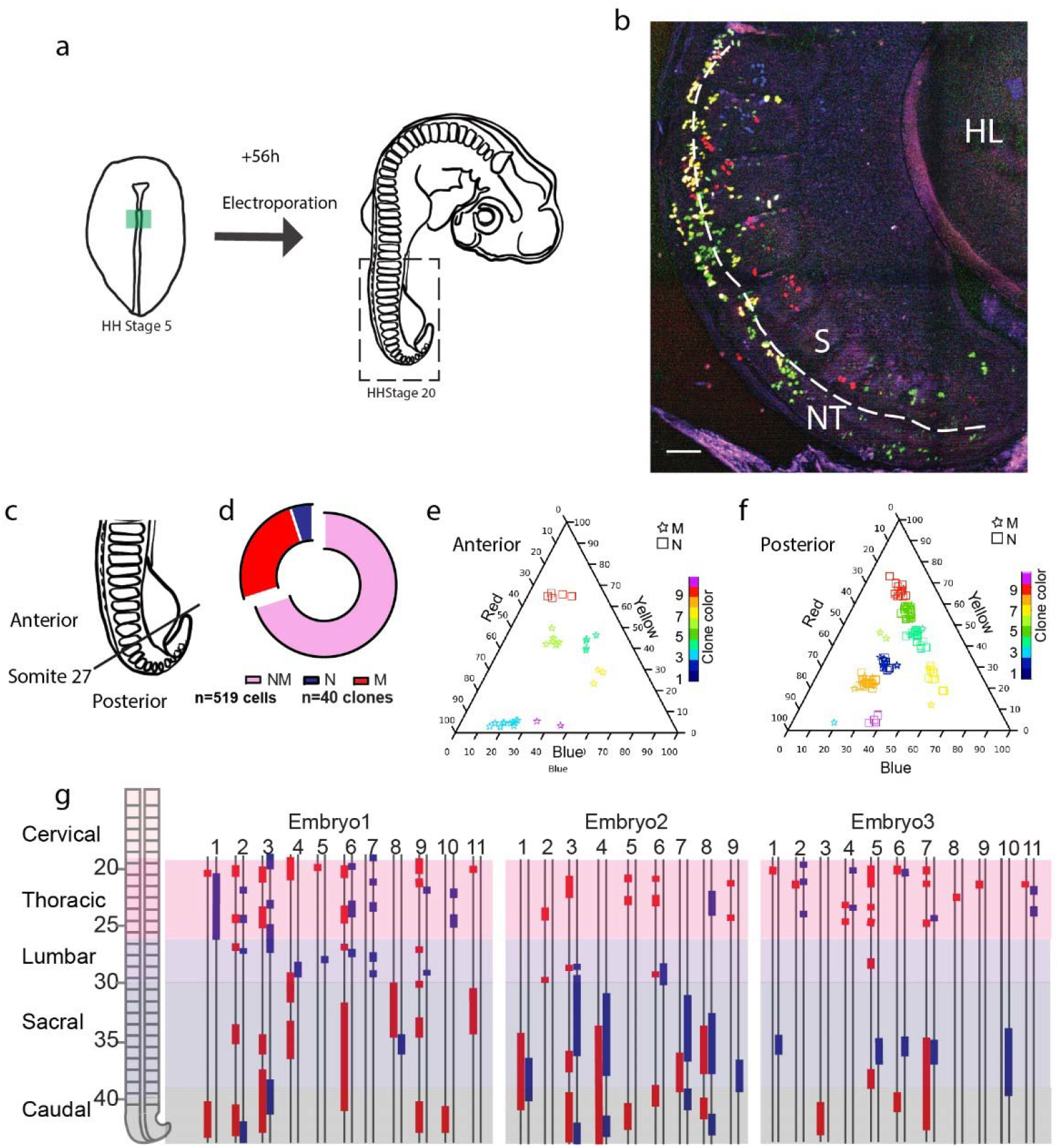
Analysis of nucbow labeled NMP clones after 72 h. (a) Experimental procedure showing the electroporated region of the epiblast at stage 5 HH (left, green) and the stage the embryos were harvested for analysis at stage 20 HH (n=4). (b) Confocal z-section using 3-color imaging (Loullier et al, 2014) corresponding to the boxed region of a stage 20 HH embryo shown in (a). (c) Definition of the anterior (above 27^th^ somite) and posterior regions (below 27^th^ somite). (d) Quantification of the different clones: Monopotent mesodermal (M,red), monopotent neural (N,blue) and bipotent Neuro-Mesodermal clones (NM, pink). (n=40 clones, 519 cells). (e-f) Triplots showing the distribution of 10 representative clones in the anterior (e) and posterior (f) region of a stage 20HH embryo electroporated at stage 5HH. Squares: neural cells, stars: mesodermal cells. (g) (Left) Region analyzed showing the different axial levels. (Right) Axial distribution of the clones in 3 embryos. Blue bars: neural cells, red bars: mesodermal cells, double line: Antero-posterior axis. Dorsal views, anterior to the top. NT: Neural Tube, S: Somite, HL: Hindlimb. Scale bar: 100μm.

**Supplementary Figure 4:**
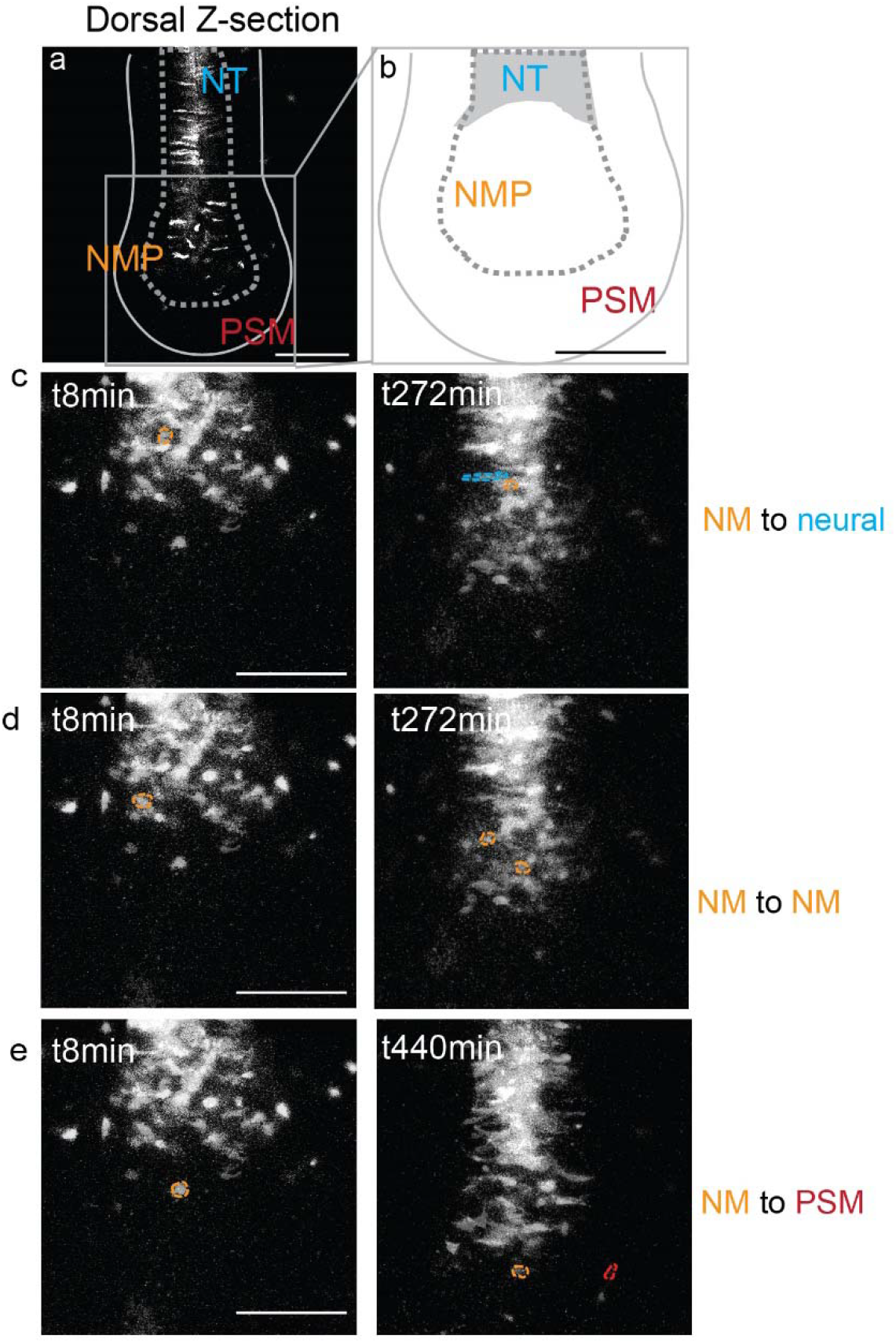
Characterization of cells of the SOX2/T region at the tailbud stage. (a) Dorsal Z section of a 25-somite embryo showing the superficial localization of the descendants from the cells co-electroporated at stage 5 HH with a GAP43-Venus and an H2B-RFP plasmid (marking the membrane and the nucleus respectively). (b) Diagram showing the dorsal region of the tail bud region where cells were tracked in c-e. (c-e) Images of tracked cells of the SOX2/T region that become neural (c), remain NMP, (d) or become PSM (e) cells. NMP cells are circled in orange, neural cells are circled in light blue and PSM cells are circled in red. (52 cells tracked in 3 embryos). NT: Neural tube, NMP: Neuro-mesodermal progenitors, PSM: Presomitic Mesoderm. Anterior to the top, dorsal views. Scale bar: 100μm

**Supplementary Figure 5:**
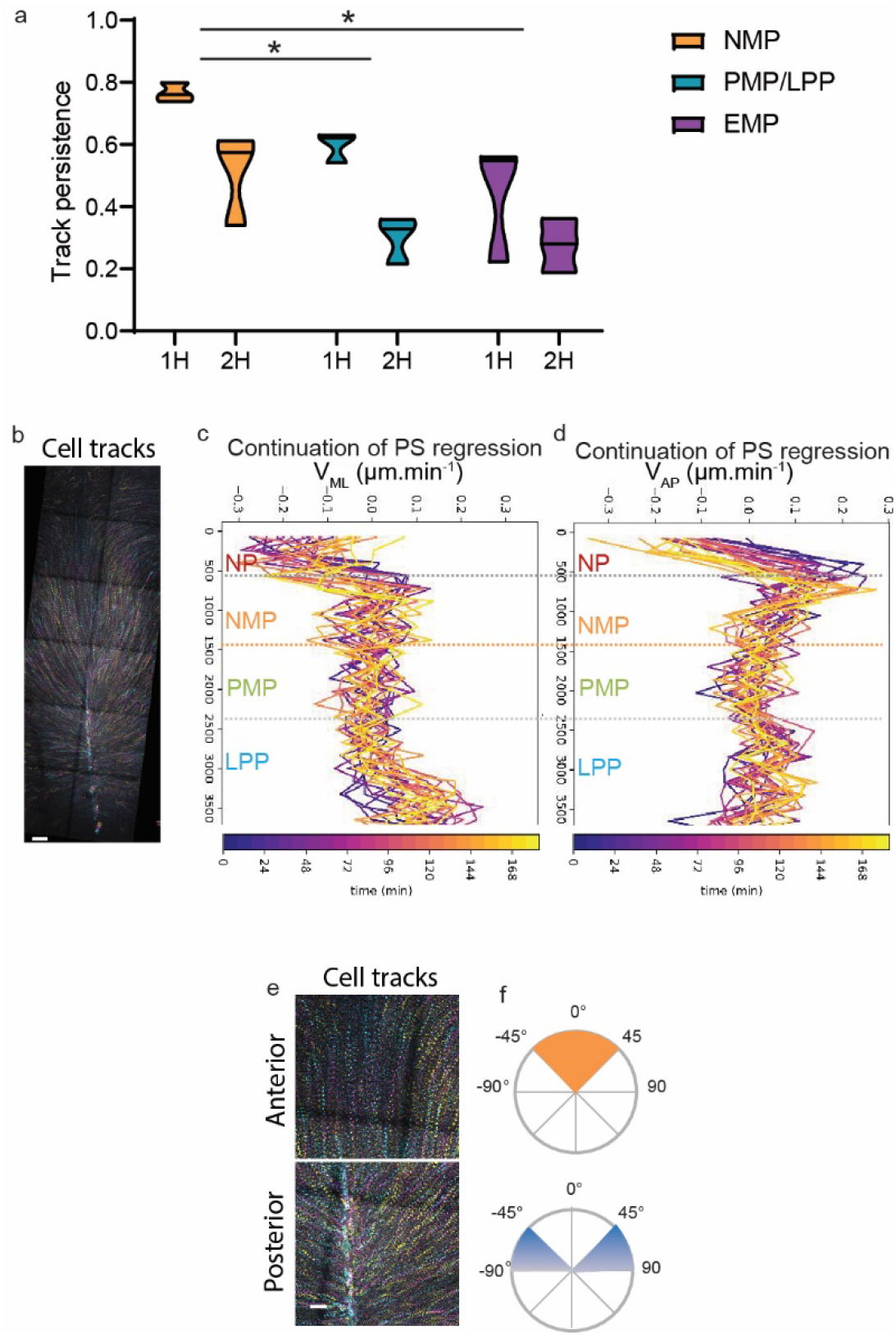
Quantification of persistence, cell speed and trajectories in time-lapse movies of nuclear red-labeled chicken embryos. (a) Quantification of tracks persistence measured as the ratio of tracks number after 1 and 2h divided by the number of tracks at t_0_ in each of the boxed regions shown in in Figure 3b. t_0_ marks the start of time-lapse movies of stage 5HH chicken embryos labeled with nuclear red. Orange, turquoise and purple show tracks in the NMP, PMP/LPP and EMP domains respectively corresponding to the tracks shown in Figure 3b. (n=3; n=1502 tracks). 2-way ANOVA NMP-PMP/LPP; NMP-EMP. *: p<0.05 (b) Maximum time color-coded projection showing the tracks of cells labeled with nuclear red in a Stage 5HH chicken embryo analyzed at the 6-somite stage. Early to later time points are indicated by cyan to magenta to yellow color code. (c-d) Mean Medial to Lateral speed (V_ML_,c) and Anterior to Posterior speed (V_AP_,d) over time of epiblast cells labeled with nuclear red in stage 5HH chicken embryos imaged by confocal imaging in vitro at the 5-somite stage. The color code indicates the time of analysis from purple to yellow where blue is t=0 minutes (stage 7 HH) and yellow is t=184 minutes. (e) Color-coded time projection of the epiblast cells labeled *in ovo* with nuclear red in stage 5HH chicken embryos, imaged by confocal imaging in vitro (e) showing different track angles with the midline quantified in (f). Early timepoints are in cyan, later timepoints are in yellow. Dorsal views, anterior to the top. NP: Neural Plate; NMP: Neuro-Mesodermal Progenitors; PMP: Presomitic Mesoderm Progenitor; LPP: Lateral Plate Progenitor. PS: Primitive Streak. (n=3 embryos). Scale bar: 100μm.

**Supplementary Figure 6:**
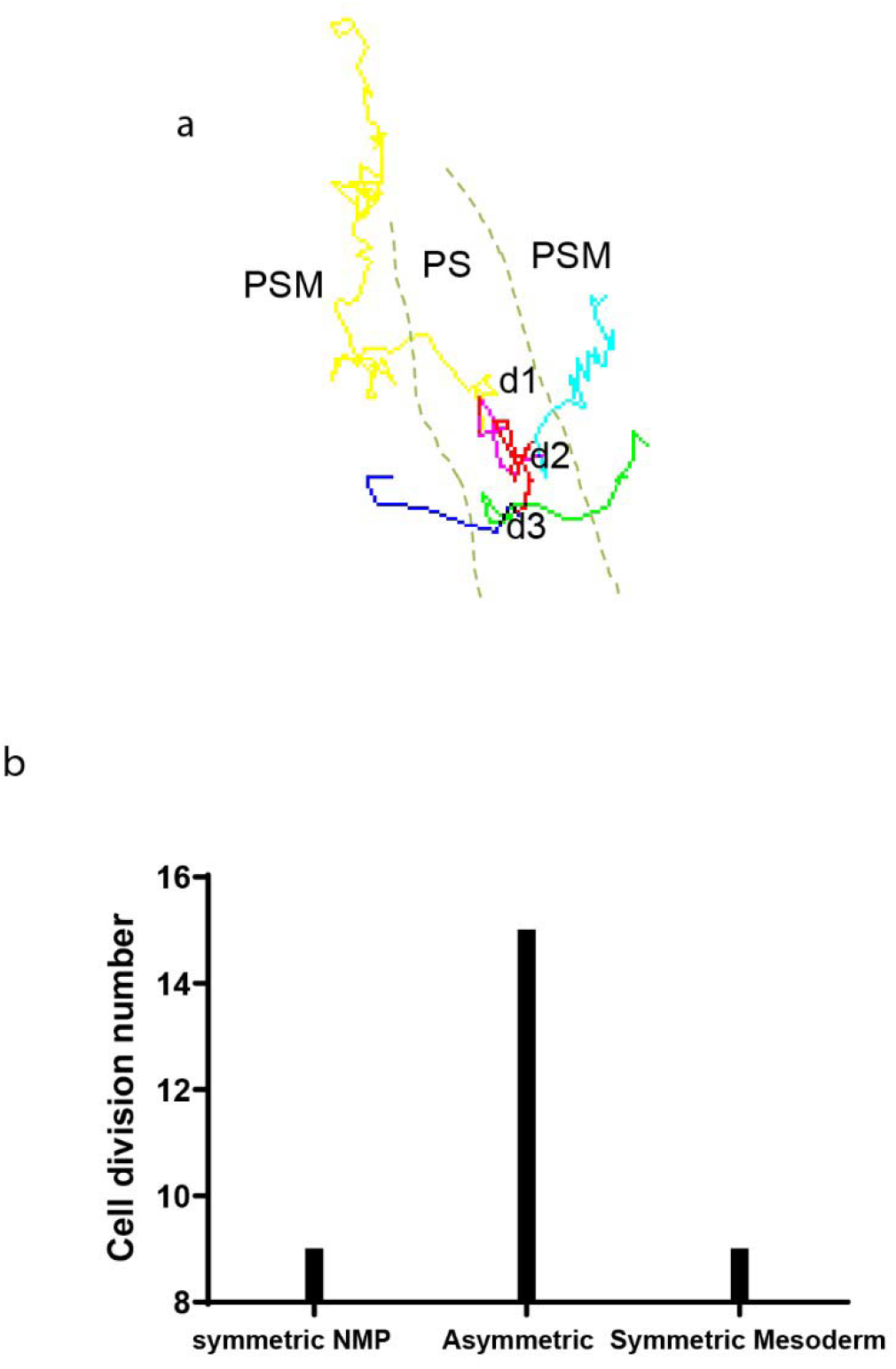
Analysis of cell division profiles in the SOX2/T region. (a) Tracks of 3 rounds of cell division of an epiblast cell of the SOX2/T positive territory in an H2B-RFP electroporated chicken embryo imaged for 12 h starting at stage 5HH (corresponding to Supplementary movie 7). d1, d2, d3: the division events. (b) Based on the location of daughter cells after cell division, we defined the cell division as symmetric (Symmetric NMP: when both daughter cells remain in the PS, Symmetric mesoderm: when both daughter cells subsequently join the mesoderm (d3)) or asymmetric (when one daughter cell remains in the PS and the other daughter cell joins the mesoderm (d1,d2). We identified 33 division events in 7 embryos which are shown in (b).

## Supplementary movies

**Supplementary Movie 1: Tracking descendants of the anterior PS epiblast at 25 somites**. Time lapse movie of dorsal maximum z-projections showing descendants from the NMP region electroporated at stage 5 HH with an H2B-RFP plasmid in time lapse. t= 4 minutes between frames, movie = 10 h. z sectioning= 4 μm. 20X objective, LSM 780.

**Supplementary Movie 2: Tracking the fate of descendants of the anterior epiblast at 25 somites**. Time lapse movie of dorsal maximum z-projections of a 25-somite embryo showing the localization and fate of descendants from cells of the anterior PS epiblast region co-electroporated at stage 5 HH with a GAP43-Venus and an H2B-RFP plasmid (marking the membrane and the nucleus respectively). The SOX2/T region of the tailbud is delimited by the white lines. Tracks of selected cells are shown. Color code indicates the z position of the cells (yellow: dorsal; blue: ventral). t=8 minutes between frames. z sectioning=4 μm 20X objective, LSM 780.

**Supplementary Movie 3: Long term tracking of a nuclear red stained embryo**. Time lapse movie of dorsal maximum z-projections of epiblast cells labeled with nuclear red showing their tracks from stage 5 HH to 5 somites during PS regression. Colors indicate the z position from magenta (dorsal) to yellow (ventral). t= 4 minutes between frames, z sectioning=4 μm. 20X objective, LSM 780. NMP: Neuro-Mesodermal Progenitors; PMP: Presomitic Mesoderm Progenitors; LPP: Lateral Plate Progenitors. PS= Primitive Streak. NT= Neural Tube.

**Supplementary Movie 4: Persistence of tracks along the PS**. 3.5 h time-lapse movie of a stage 5HH chicken embryo in which the epiblast was labeled with nuclear red. Tracks of single labeled nuclei are shown at three different AP levels of the PS to illustrate differences in persistence of the cells. Dorsal view. (NMP: orange; PMP/LPP: green; EMP: blue).t=4 minutes 20X objective, LSM 780.

**Supplementary Movie 5: Dynamics of mesodermal cell ingression**. Time lapse movie showing ventral z-maximum projections of the PS region of a chicken embryo labeled dorsally at stage 5HH with nuclear red. t=15 min z=4 μm 20X objective, LSM 780.

**Supplementary Movie 6: Long term tracking of the mesodermal progenitors**. 13h time lapse movie of a GFP chicken embryo starting at stage 5HH (Left). Fate of the color-coded mesodermal PS progenitor territories during regression is shown in the right movie. Neuro-Mesodermal Progenitors: NMP, orange; Presomitic Mesoderm Progenitors: PMP, Green; Lateral Plate Progenitor: LPP, Extra-embryonic Mesoderm Precursors, LPP-EMP, Blue-purple. Arrowhead: Hensen’s Node. Anterior to the top. t=4 min, 20X objective, LSM 780.

**Supplementary Movie 7: Tracking cell division time in the SOX2/T region.** Tracking of cell divisions in cells of the SOX2/T region electroporated with an H2B-RFP plasmid at stage 5 HH. d1, d2, d3: division events, PS: Primitive Streak. t=6 minutes. 10X objective Leica DMR.

## Material and methods

### Chicken embryos

All animal experiments were performed in accordance to all relevant guidelines and regulations. The office for protection from Research Risks (OPRR) has interpreted “live vertebrate animal” to apply to avians (e.g., chick embryos) only after hatching. All of the studies proposed in this project only concern early developmental stages (prior to 5 days of incubation), therefore no IACUC approved protocol is required. Fertilized chicken eggs were obtained from commercial sources. Fertilized eggs from transgenic chickens expressing cytoplasmic GFP ubiquitously (McGrew et al., 2004) were obtained from Susan Chapman at Clemson University. Fertilized eggs from transgenic quails expressing H2B-Cherry (Benazeraf et al., 2017) were obtained from Ozark Hatcheries. Eggs were incubated at 38 °C in a humidified incubator, and embryos were staged according to Hamburger and Hamilton (HH) (Hamburger and Hamilton, 1992). We cultured chicken embryos mainly from stage 5HH on a ring of whatman paper on agar plates as described in the EC culture protocol (Chapman et al., 2001).

### Immunohistochemistry

For whole mount immuno-histochemistry, stage 3 to 20 HH chicken embryos were fixed in 4% paraformaldehyde (PFA)(158127,Sigma) diluted in PBS 1X at 4 °C overnight. The embryos were rinsed and permeabilized in PBS-0.1% triton, 3 times 30 min, and incubated in blocking solution (PBS-0.1% triton, 1% donkey serum (D9663, Sigma)) prior to incubating with primary and secondary antibodies. Embryos were incubated in antibodies against T/BRACHYURY (1/1000, R&D Systems: AF2085), SOX2 (1/1000, Millipore: ab5603), E-CADHERIN (1/250 Abcam: ab76055), N-CADHERIN (1/250, Abcam: ab12221), FIBRONECTIN (1/50 DSHB: MT4S), LAMININ (1/200, SIGMA: L9393), PHOSPHO-HISTONE 3 (1/1000, SANTA CRUZ: sc-8656), MSGN1 (1/1000) (Oginuma et al., 2017), diluted in blocking solution at 4 °C overnight. Embryos were rinsed and washed 3 times 30 minutes in PBS-0.1% triton, incubated 1h in blocking solution and incubated at 4 °C overnight with secondary antibodies conjugated with AlexaFluor (Molecular probes) diluted in blocking solution. If the staining was not imaged in the following 2 days, post fixing was performed using a 4% PFA solution.

For histological analysis, stage 5 HH chicken embryos were fixed in 4% PFA. Embryos were then embedded in OCT compound and frozen in liquid nitrogen. Frozen sections (12 μm) were cut using a Leica Cryostat and incubated overnight at 4 °C with the primary antibody diluted in blocking solution (same as above), and after washing in PBS-0.1% triton, they were incubated overnight at 4 °C with the secondary antibody conjugated with Alexa Fluor (Molecular probes) diluted in blocking solution. Sections were then washed in PBS-0.1% triton before mounting in fluoromount-G (thermofisher) and stored at 4°C overnight prior imaging.

Images were captured using a laser scanning confocal microscope with a 10X or 20X objective (LSM 780, Zeiss). To image the whole embryo, we used the tiling and stitching function of the microscope (5 by 2 matrix) and z sectioning (5μm). Later stages (from HH17) were imaged in clearing solution using the scale A2 clearing protocol from (Hama et al., 2011). For imaging, the embryo was placed in the clearing solution (4 M urea 0.1% Triton, 10% glycerol) 30 minutes prior imaging in glass bottom dishes (Mattek).

### Plasmid preparation and electroporation

pCAGG-H2B-Venus and pCAGG-H2B-RFP have been described in (Denans et al., 2015), pCAGG-GAP43-Venus has been described in (Oginuma et al, 2017). Chicken embryos at stage 4 or 5HH were prepared for in ovo electroporation. Eggs were windowed and a DNA solution (1 μg/μl) mixed in PBS, 30% glucose and 0.1% Fast-green was microinjected in the egg, in the space between the vitelline membrane and the epiblast in the first 500 μm posterior to the node or 1500 μm posterior to the node to target the NMP or the LP progenitors respectively. Electroporation was carried out using 2 pulses at 5V for 1msec on each side of the PS in the NMP and LP domains (4 locations) using a needle electrode (CUY614, Nepa Gene, Japan) and an ECM 830 electroporator (BTX Harvard Apparatus). This procedure only labels the superficial epiblast layer (Iimura and Pourquie, 2008). Eggs were then re-incubated for 1 to 2 hours at 38°C and embryos were dissected and prepared for EC culture imaging in 6 well imaging plates as described in (Denans et al., 2015). For lineage tracing experiments, we performed the same procedure as above but using only 1 pulse on each side of the PS in the NMP domain to minimize the number of electroporated cells.

### Nuclear red labelling in vivo

The nuclear red solution was prepared from the NucRed™ Live 647 ReadyProbes™ Reagent (Thermofisher) and diluted in PBS1X as indicated by the manufacturer experimental procedures. Sparse nuclear labelling of the dorsal epiblast was performed in ovo by injecting the nuclear red solution between the epiblast and the vitelline membrane at the PS level. The solution was left for 15 minutes to perform the long-term epiblast tracking and 30 minutes for monitoring ingression. The embryos were then dissected, rinsed in PBS and mounted on paper filter for EC culture to perform live imaging from the dorsal side for the long-term epiblast tracking and from the ventral side for measuring cell ingression.

### Time-lapse imaging, ingression and cell cycle length measurements

Stage 5HH chicken embryos were cultured ventral side up on a microscope stage using a custom built time-lapse station (Benazeraf et al., 2010). We used a computer controlled, wide-field (10× objective) epifluorescent microscope (Leica DMR) workstation, equipped with a motorized stage and cooled digital camera (QImaging Retiga 1300i), to acquire 12-bit grayscale intensity images (492 × 652 pixels). For each embryo, several images corresponding to different focal planes and different fields were captured at each single time-point (frame). The acquisition rate used was 10 frames per hour (6 min between frames). To quantify cell ingression and division, the image sequence was registered to the node displacement by tracking the advancement of the Hensen’s Node as a function of time using the manual tracking plug-in in Image J (Denans et al., 2015). Cell division events were manually tracked in long-term movies of H2B:GFP and RFP electroporated embryos. The time between divisions was estimated by counting the number of frames.

To analyze the dynamics of mesodermal precursor territories during PS regression, we tracked three small regions of the epiblast adjacent to the PS located at a distance of 500, 700 and 1000 μm from Hensen’s node at stage 5HH in transgenic chicken embryos expressing GFP. Embryos were imaged with a zeiss LSM 780 confocal microscope with temperature control. At stage 5HH, the entire PS is around 2000 μm in length. According to our fate mapping (see Figure 1) and to Psychoyos and Stern (1996), these three regions correspond to transition zones between the NMP-PSM, PSM-LP and LP-EM territories respectively. The NMP territory is maintained over the first 500 μm from the node and remains in the *sinus rhomboidalis* region during PS regression. The PSM progenitor domain is found posterior to the NMP territory in the anterior third of the PS approximately extending 670 μm posterior to the Node. We observed that electroporation of the region located 750 μm posterior to the Node is labelling LP cells (Figure 3). Thus, we tracked the area located 700 μm posterior to the Node which approximately corresponds to the transition zone between the precursors of paraxial mesoderm and LP. The posterior half of the PS in stage 5HH chicken embryos mostly gives rise to extra-embryonic mesoderm (Psychoyos and Stern, 1996). Thus, we considered the boundary between lateral plate and extra embryonic mesodermal progenitors to be at 1000 μm from Hensen’s Node. These different regions were followed using the manual tracking module from imageJ in 10 h time-lapse movies spanning from stage 5HH to the 10-somite stage. Supplementary movie 6 shows a representative movie where the different territories have been color-coded to visualize their dynamics during PS regression.

### Cell tracking analysis

Whole epiblast cell tracking was performed using the MOSAIC plugin from Image J (Sbalzarini and Koumoutsakos, 2005). Tracks were then visualized and analyzed using a custom code in Python. Cell velocities were computed by calculating the discrete displacements. In order to back-track cells in regions of interest, groups of cells were selected at any given time using selection tools provided by the Scikit-image package (van der Walt et al., 2014). Trajectories were then plotted using the Matplotlib package (Hunter, 2007). Color-coded time projections were generated using the color-coded projection module from Zen software. Early to late timepoints are shown using the following color sequence: cyan, magenta and yellow.

### Cell proliferation analysis

Stage 5, 8,12,13 HH chicken embryos were dissected, fixed in 4% PFA and immunostained with Phospho-histone H3 (pH3) antibody (1/1000, Millipore) as described above. In the penultimate wash, Hoechst (1/1000) was added to label cell nuclei. The number of proliferative cells was measured in 250 by 250 μm squares at 3 different locations along the PS (center of the area being at 250, 750, 1200μm from the node). Proliferative cells were identified by automated segmentation of pH3 positive cells using ImageJ via tresholding the images and using the analyze particle module. Hoechst-positive cells were also segmented using the same procedure and their number was used to normalize to the cell density in each square. The percentage of dividing cells was obtained by dividing the number of pH3 cells by the number of Hoechst positive cells in each square and multiplying by 100. A minimum of 5 embryos was analyzed for each stage. We also performed similar measurements on live H2B: Cherry quail embryo at stage 5HH in the anterior (around the node) and posterior (Lateral plate progenitor region). We used a high threshold, which identifies cells in mitosis (that show a brighter cherry signal due to chromatin condensation), while all the other cells in the tissue are identified with a lower treshold.

### Analysis of the localization and number of NMPs

Chicken embryos from stage 4 to 20 HH were harvested and fixed in 4% PFA at 4C overnight. Embryos were immunostained in whole mount using T/BRACHYURY, SOX2 and Alexa Fluor™ 647 Phalloidin (1/500 Thermo Fisher: A22287) combinations and imaged by confocal laser microscopy (LSM 780, Zeiss). SOX2/T double positive cells were segmented and counted using image J software. To identify the double positive cells, cells in the T and in the SOX2 channel were thresholded. We used the image calculator tool from ImageJ, to generate a new image by performing the image operation T AND SOX2 and identify the double-positive cells. Embryos were grouped by stage and data is shown in Figure 4a. n=7 at stage 4HH; n=11 at stage 5HH; n=6 at 5-10 somites; n=5 at 15-20 somites; n=5 at 30 somites; n=4 at 35 somites and older. Total n=38 embryos.

### Lineage tracing and quantification

#### Cloning of a High Complexity Barcoded Retroviral Genomic Plasmid Library

The plasmid library used to generate the barcoded retrovirus was based on the pQCXIX backbone. First, this backbone was digested with NotI and XhoI. A Gibson assembly reaction was performed using three PCR generated fragments to insert the EGFP, IRES, and Cre Recombinase, creating pQCGIC (Gibson et al., 2009). These PCR fragments were generated using the indicated primers from pCAG-EGFP, pQCXIX, and pCAG-Cre plasmids, respectively. This plasmid was then digested with XhoI and PvuII and an intron disrupted polyadenylation sequence (IDPA) amplified from a gBlock (IDT) was inserted with a Gibson assembly reaction, creating pQCGICIDPA. Finally, the dsDNA barcode insert sequence was prepared from an Ultramer Oligo (IDT), QCGICbc_oligo, with PCR. This ultramer was synthesized with 12 SW repeats, resulting in a GC balanced 24 bp barcode. pQCGICIDPA was digested with XhoI, and a 170 μl Gibson assembly reaction was performed with 8.5 μg of linearized vector and 1.24 μg of insert. This reaction was purified and electroporated into three 100 μl aliquots of electrocompetent DH10β *e. coli*. Following a 1 h outgrowth in 12 ml SOC at 37C with shaking, dilutions were plated on LB agarose plates with carbenicillin to determine complexity, and the remaining transformation was added to 500 ml of LB with carbenicillin and grown for 16 hours at 37C. The liquid culture was split into 4 and plasmid was purified with the Qiagen Maxiprep Kit. A plate with 1/1×10^4^ of the transformation grew approximately 1×10^3^ colonies, indicating a total library complexity of 1×10^7^.

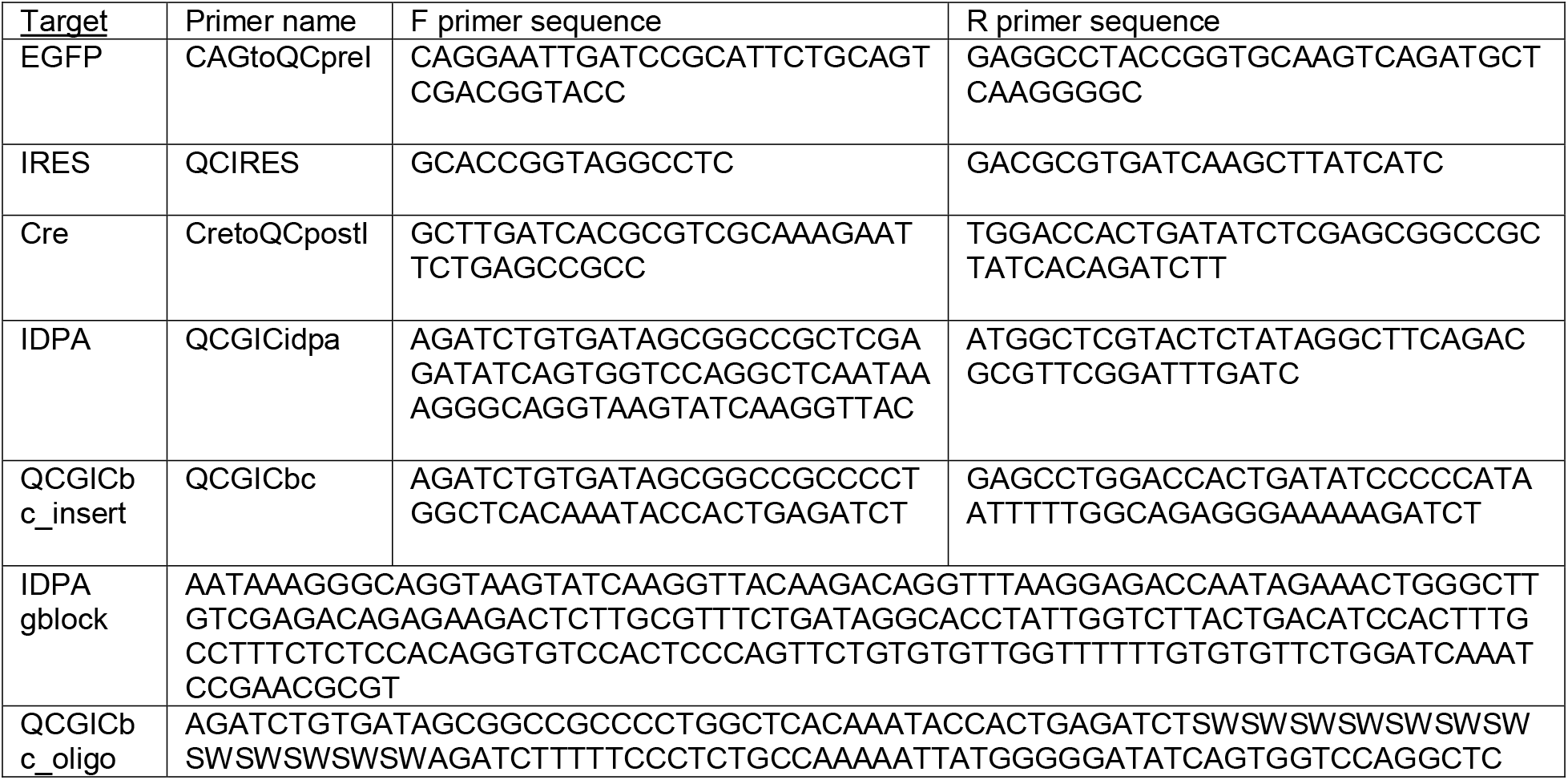

#### Preparation of Replication-Incompetent Retrovirus

Replication incompetent retrovirus was produced and concentrated as previously described (Beier et al., 2011; Cepko and Pear, 2001). Briefly, on Day 1, eleven 150 cm plates with HEK 293T cells at 70%-80% confluency were transiently transfected each with 13.5 μg of pQCGICbcIDPA, 6.75 μg pMN gagpol (Ory et al., 1996) and 2.25 μg of a plasmid expressing the EnvA envelope (Landau and Littman, 1992) using PEI (polysciences, 24765; 90 μg per plate). Media was replaced on Day 2 and viral supernatant was collected on Day 3 and Day 4 and stored at −80C. The supernatant was thawed, and virus was concentrated via ultracentrifugation (49,000 xg for 2 hours at 4C). Virus was resuspended in 270 μl of media and aliquoted in 10-20 μl aliquots and stored at −80C. Titer was determined by infecting HEK 293T cells expressing the avian TVA receptor and was found to be approximately 4×10^8^ infectious particles per milliliter.

#### Retroviral infection

1μl of the virus preparation was diluted in 10 μl of PBS1X and 0.1% fast green to form the virus mix. The virus mix was placed in ovo on top of the NMP region for 1 hour and washed with PBS 1X. The eggs where then sealed with tape and reincubated for 36 h.

#### Quantification

To retrieve the different barcodes, we manually harvested individual fluorescent cells from transverse sections of embryos fixed 36 h post-infection. GFP-positive cells were handpicked under a microscope (20X objective) using single use needles (BD precision). Each cell/needle was put in an individual PCR tube containing the buffer for the PCR1 and its localization was recorded. The tissue of origin of the cells was visually identified on the sections. The barcodes were retrieved using 2 consecutive nested PCR amplifications and sequenced using sanger sequencing. PCR primer 1: 1R-TCTCTGTCTCGACAAGCCCAG, 1F-GATCATGCAAGCTGGTGGCTG. PCR primer 2: 2R-CTTACCTGCCCTTTATTGAGCCTG, 2F-CTGCTGGAAGATGGCGATGG.

#### Nucbow cell tracing

Lineage tracing was performed by co-electroporating the NMP region at stage 5HH *in ovo* as described above with the following constructs: a self-excising Cre recombinase (se-Cre), the nucbow construct and the TolII transposase as described in Loulier et al (2014) in a 1/1/1 ratio at (1μg/μl, each). We used similar concentrations for the nucbow and transposase plasmids to that described in Loulier et al, (2014) but increased 10 times the concentration (1 μg/μl versus 0.1 μg/μl) of the se-Cre to favor fast recombination and integration. Because non-integrated nucbow plasmids can remain episomal and transiently affect the color of a cell, we performed our analyses after 36 h when the plasmids are expected to have fully diluted through cell division. 36 h after electroporation, we see that the number of fluorescent cells has significantly decreased suggesting that the episomal transgenes have now been diluted.

To perform lineage analysis, we fixed the electroporated embryos at stage 17HH and 20HH, and imaged them in clearing solution ScaleA2 (Hama et al., 2011). The imaging was performed using an LSM 880 with Airyscan module in the 3 fluorescent channels following the same gating as in (Loulier et al., 2014).

#### Quantification

Cells were manually segmented in the YFP and Cherry channel using image J. Positions were assigned to the mesodermal and neural tube layers. Color retrieval was performed by measuring the intensity in the 3 channels, Cerulean, YFP and Cherry so that the total of all the intensities was normalized to 1 and expressed in percentages similarly to (Loulier et al., 2014). Cluster assignment was performed using K-mean clustering followed by thresholding of only the cells with a silhouette >0.4. The coordinates were then calculated in a triplot diagram for visualization. The cells in a clone were then re-assigned to their original localization based on the Y value as a proxy for the Antero-posterior position in the image.

